# Single-Session Fast-Paced Video Gaming Boosts Working-Memory Precision via Prefrontal Alpha Oscillations

**DOI:** 10.1101/2025.01.03.631246

**Authors:** Ali Asgari, Abdol-Hossein Vahabie, Ehsan Rezayat

**Affiliations:** Department of Cognitive Sciences, Faculty of Psychology and Education, University of Tehran, Tehran, Iran; School of Electrical and Computer Engineering, College of Engineering, University of Tehran, Tehran, Iran; School of Cognitive Sciences, Institute for Research in Fundamental Sciences (IPM), Niavaran, Tehran, Iran

**Keywords:** Working-memory precision, High-alpha oscillations, Dorsolateral prefrontal cortex, Fast-paced video game, Resting-state EEG, Cognitive plasticity

## Abstract

Working-memory (WM) precision depends on top-down control by dorsolateral prefrontal cortex (dlPFC). Whether a brief session of action video-game play can sharpen this control, and which neural processes mediate any benefit, remains unknown. Thirty-four adults were randomly assigned to 30 min of fast-paced gaming (Snail Mail; Experiment) or time-matched cartoon viewing (Control). Delayed continuous-report precision and 128-channel resting EEG were collected before and after the intervention. Absolute recall error fell in the Experiment group (p < .001) but rose in Controls (p = .005); the groups did not differ at baseline (p = .164) yet the Experiment group outperformed Control at post-test (p < .001). In the EEG subset (N = 30), gaming selectively increased high-alpha (10–13 Hz) power over left dlPFC (AFF7h; p = .041), an effect absents in Controls and significant between groups (p = .010). Across participants, the magnitude of this prefrontal alpha gain predicted the improvement in WM precision (r = .53, p = .0025), whereas concurrent flattening of the global 1/f slope was unrelated (|r| < .24, p > .21). A single, ecologically realistic gaming session therefore yields measurable enhancement of mnemonic precision together with local up-regulation of dlPFC-centered alpha oscillations, identifying a plausible cortical mechanism for rapid, low-cost cognitive plasticity.

## Introduction

Working-memory (WM) is the executive workspace that allows humans to maintain and manipulate information in the service of current goals. Contemporary frameworks converge on a multicomponent architecture in which dedicated buffers interact with an executive controller (Baddeley 2000). Converging lesion, neurophysiological and neuro-imaging data localize the highest level of that control to dorsolateral prefrontal cortex (dlPFC), a region necessary for the manipulation, but not the sheer maintenance, of stored representations (Barbey, Koenigs & Grafman 2013). Classic integrative theory proposes that dlPFC accomplishes this by maintaining goal-related activity patterns that bias processing in posterior cortices (Miller & Cohen 2001), a view refined by more recent evidence for distributed, population-level coding across PFC–sensory networks (Sreenivasan et al. 2014). Behaviorally, WM precision, rather than simple capacity, is the parameter most sensitive to such top-down control (Bays, Catalao & Husain 2009).

A large electrophysiological literature suggests that oscillations in the α-band (8–13 Hz) are instrumental in this control. Alpha power increases at task-relevant cortical sites are thought to gate processing by actively inhibiting competing input (Klimesch 2012), thereby protecting WM representations from interference (Bonnefond & Jensen 2012). In visuospatial WM, α power typically rises over prefrontal leads while concurrently decreasing over occipital cortex, signaling targeted suppression of sensory influx (Sauseng et al. 2005). Upper-α (≈10–12 Hz) power at rest predicts individual memory performance (Vogt, Klimesch & Doppelmayr 1998) and can be up-regulated through neurofeedback to enhance cognition (Zoefel, Huster & Herrmann 2011). Despite these links, it remains unclear whether brief, ecologically valid experiences can causally modulate frontal α activity and whether such modulation translates into immediate gains in WM precision.

Action video games constitute a promising candidate intervention. Habitual players outperform non-players across a range of visual–attentional measures (Green & Bavelier 2003), and longitudinal training studies demonstrate transfer to untrained cognitive domains, including top-down control, in both young and older adults (Anguera et al. 2013; Bediou et al. 2018). The mechanisms, however, are debated: some authors attribute transfer to improved learning efficiency (Green & Bavelier 2012), whereas others question the generalizability of game-based “brain training” (Simons et al. 2016; von Bastian et al. 2022). Electrophysiological evidence is sparse, and few studies have examined whether a single session of fast-paced gaming can elicit measurable neural plasticity relative to a non-interactive control such as passive cartoon viewing (Fan et al. 2022).

Although months-long interventions show that action gaming can reshape perceptual attention, the *timescale* and *neural locus* of such plasticity remain poorly understood. Most randomized trials employ ≥ 10 h of training spread over weeks (Bediou et al., 2018) and assess broad cognitive composites rather than fine-grained WM parameters. Consequently, it is unclear whether the key behavioral benefit, *enhanced mnemonic precision*, emerges only after extensive practice or can already be detected after a single, ecologically realistic bout of game play.

At the neural level, three gaps stand out. First, EEG studies of game interventions have so far focused on posterior visual rhythms linked to vigilance, leaving the putative executive generator in dlPFC largely unexplored. Second, the frequency specificity of gaming-induced changes is unresolved: while long-term gamers sometimes show reduced *low-α* power (≈ 8–10 Hz) over parietal cortex, theoretical models predict that task-engaged *high-α* in prefrontal regions should *increase* to shield WM from distraction (Klimesch, 2012; Bonnefond & Jensen, 2012). Third, recent work highlights the importance of disentangling periodic α peaks from the broadband aperiodic (1/f^χ) background that indexes cortical excitation–inhibition balance (Donoghue et al., 2020). No study has yet asked whether rapid game play alters χ and, crucially, whether any such change relates to behavioral precision.

The present experiment addresses these issues with a tightly controlled, pre-registered design. We contrasted 30 min of *Snail Mail*, a commercially available, fast-paced arcade game relying exclusively on mouse steering, with a matched period of passive cartoon viewing. Both activities are engaging (Boyle et al., 2012) yet differ fundamentally in interactive demands, providing an ideal test of causal effects. We assessed WM precision immediately before and after the intervention and recorded high-density (128-channel) resting EEG after the task; quantified first, ΔWM precision, pT, and pU from a three-face delayed-reproduction task. Second, ΔHigh-α relative power at a pre-defined left-dlPFC ROI (AFF7h) and across the scalp; Third, ΔAperiodic slope (χ) and offset, extracted with both robust log–log fits and FOOOF.

### Pre-registered hypotheses

#### Behavior

The Experiment (gaming) group will exhibit a larger post-minus-pre reduction in absolute WM error (i.e., increased precision) than the Control (cartoon) group. *EEG*: Gaming will selectively increase high-α power over dlPFC, with no comparable change in Control. *Brain– behavior link*: Across participants, ΔHigh-α at AFF7h EEG channel (dlPFC) will positively correlate with ΔPrecision, supporting an α-mediated top-down mechanism. *Exploratory*: We probe whether gaming flattens the aperiodic slope (less negative χ) and whether χ relates to precision after controlling for group.

### Single-session plasticity as a stringent test of causality

Most action-game studies aggregate tens of training hours, leaving open whether benefits accrue gradually or whether an early, short-lived “boost” drives later gains. Demonstrations of rapid neuromodulation would strengthen causal inference by minimizing confounds such as differential attrition and clarifying the temporal dynamics of cognitive plasticity (von Bastian et al., 2022). By sampling resting EEG immediately before and after the working memory task, we target neural changes on the order of minutes, an interval in which long-term potentiation–like processes at fronto-cortical synapses are plausible.

### An active, content-matched control

Passive video viewing was selected because it matches the visual flow and motivational appeal of gaming (Boyle et al., 2012), and brief exposure to entertainment cartoons can *impair* executive functions in children (Fan et al., 2022) and adults, presumably by promoting externally oriented attention. Any advantage of gaming over cartoons therefore reflects the added requirement to generate rapid predictions, action plans and error corrections rather than banal sensory stimulation. Importantly, both interventions were administered in the same seated laboratory context, with identical experimenter contact and timing, satisfying consort guidelines for behavioral trials.

### A precision-weight WM paradigm that taxes prefrontal control

Delayed reproduction of a continuous face-emotion ring isolates mnemonic precision from guess rates and has revealed selective dlPFC engagement even when storage load is equated (Bays et al., 2009). Because precision is continuously distributed, small improvements are statistically detectable with modest sample sizes while remaining behaviorally meaningful (≈ 5–10 % reduction in mean error).

### Predicted neural signatures

We focus on *high-α* (10–13 Hz) for two reasons. First, preparatory increases in this sub-band over task-relevant cortex protect WM against anticipated distraction (Bonnefond & Jensen, 2012) and scale with individual precision (Vogt et al., 1998). Second, fast-paced gaming imposes sustained top-down filtering of irrelevant optic flow, a demand expected to up-regulate frontal α while suppressing occipital α (Sauseng et al., 2005; Mazaheri et al., 2014). We therefore predict a fronto-central, left-lateralized gain in high-α power — centered on AFF7h/5h/3h — accompanied by a relative decrease over occipital sites, reflecting a transient re-balancing of cortical excitation. The inclusion of aperiodic slope/offset metrics allows us to test whether any α change merely mirrors broadband shifts in excitation–inhibition or constitutes a frequency-specific modulation.

By combining a state-of-the-art continuous-report paradigm with high-density EEG, the present study aims to move the debate on video-game transfer beyond broad behavioral indices toward mechanistic insight. Three innovations merit emphasis. First, we focus on *mnemonic precision* instead of coarse accuracy or span, providing a sensitive assay of dlPFC-mediated control (Bays et al., 2009). Second, we dissect the neural spectrum into *periodic* (oscillatory) and *aperiodic* components. The latter, indexed by the 1/f slope, tracks global shifts in cortical excitation– inhibition balance (Donoghue et al., 2020) and therefore offers a complementary window onto the physiological impact of gaming that has been largely ignored in prior training work. Third, we pit a fast-action game against a naturalistic, equally engaging but non-interactive cartoon control, thereby isolating the contribution of active sensorimotor demands while controlling for arousal and screen time (Boyle et al., 2012).

We preregistered three predictions: *(i)* A single 30-min bout of action gaming will selectively *improve WM precision* relative to cartoon viewing. *(ii)* This gain will be paralleled by an *increase in post-intervention high-α (10–13 Hz) relative power* over left dlPFC (AFF7h), reflecting enhanced top-down gating. *(iii)* Across individuals, *larger α increases will predict larger precision gains*, attesting to a functional link between the neural and behavioral plasticity. Exploratory analyses will test whether changes in the aperiodic slope mediate or modulate these effects.

Establishing such rapid, coupled plasticity would strengthen the case that action games can transiently tune neural circuits underlying executive control, offering a laboratory-tractable model of experience-dependent optimization. Conversely, null or control-favored outcomes would temper enthusiasm for single-session interventions and clarify boundary conditions for cognitive transfer. The findings promise to refine theoretical accounts of WM regulation, α-band function and the malleability of cognitive control in adulthood.

## Method

### Ethics and participants

After a complete verbal description of the study, all participants provided written informed consent before participating and were free to withdraw at any time. All procedures were carried out in accordance with relevant guidelines and regulations; procedures were in accordance with the Declaration of Helsinki and approved by local ethics committees, i.e. University of Tehran Science Committee (IR.UT.PSYEDU.REC.1403.019).

Thirty-six right-handed adults (18 women, 18 men; 18–34 years, *M* = 23.8, *SD* = 3.8) with normal or corrected-to-normal vision and no self-reported neurological, psychiatric history were recruited from campus advertisements. Pre-test exclusion: 2 participants whose absolute-error distribution contained > 3 SD outliers were removed → *N* = 34 for behavioral analyses (Experiment = 17, Control = 17). EEG-quality exclusion: 4 further datasets (2 per group) were discarded because artefact-subspace reconstruction (ASR) removed > 25 % of samples; their behavioral data were retained, leaving *N* = 30 complete EEG datasets (15 per group).

### Experimental design

Figure 1a summarizes the ∼2 h within-session protocol. Screen position, chair height and ambient lighting were identical in both conditions. Participants were instructed to minimize head/trunk movement and refrain from talking while recording. After electrode preparation (≈ 30 min) each participant completed Pre-test working-memory task and Resting-state EEG, 7 min eyes-open (EO) followed immediately by 7 min eyes-closed (EC) while seated upright.

**FIGURE 1.**
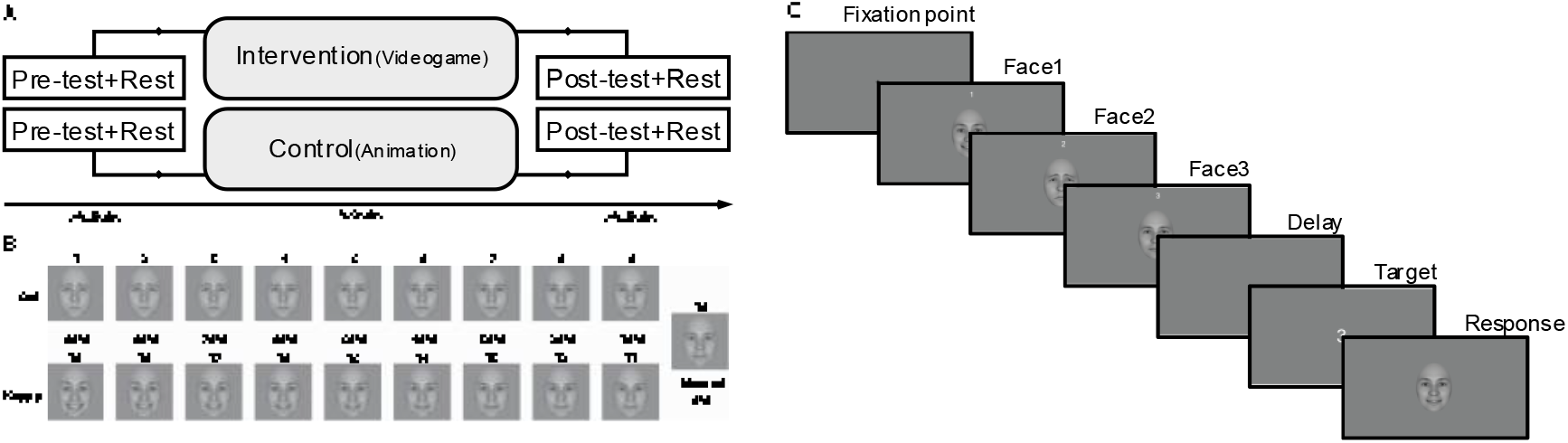
Experimental timeline and working-memory (WM) task *(A)* Two-hour session structure: baseline WM task (3 × 30 trials) → 7-min resting EEG → 30-min intervention (fast-paced game Snail Mail for the Experiment group, cartoon viewing for the Control group) → post-intervention WM task → post-rest EEG. *(B)* Nineteen morphs along a sad-to-happy continuum (10 % steps; examples shown every 20 %). Morphs 2, 6, 10, 14, 18 served as sample faces. *(C)* Delayed continuous-report trial. After a 0.7 s fixation dot, three faces (0.7 s each) were presented with serial numerals. Following a 1.5 s or 3 s retention interval, a cue numeral indicated which face to reproduce. Participants scrolled the 19-item ring with ↑/↓ keys and confirmed with *Enter* (≤ 20 s).

Participants were then randomly assigned (sealed-envelope, 1 : 1) to *Experiment group*, 30 min of the arcade-action game *Snail Mail* (Sandlot Games, 2004; Challenge mode). The game was played entirely with a standard mouse; difficulty and speed rose by 20 % steps after each stage. Or randomly assigned to *Control group*, 30 min of non-verbal cartoons (*Shaun the Sheep*, Aardman Animations). Four ≈ 7 min episodes were shown; pauses between episodes matched the stage-loading breaks in the game. Immediately afterwards they repeated the WM task (post-test) followed by an identical EO → EC resting sequence. The EO-first order reproduces typical fatigue trajectories and aligns with earlier short-term gaming studies. Total session duration, including consent and debriefing, was ≈ 120 min.

### Working-memory task

A gender-neutral base face (FaceGen Modeller 3.5) was morphed along a sadness to happiness axis in 10 % steps using InterFace 1.1, yielding 19 greyscale bitmaps (400 × 572 px) numbered 1 (90 % sad) … 19 (90 % happy). After oval cropping and luminance matching (SHINE 2.0), each face subtended 3.33° × 4.76° at 60 cm. Stimuli were presented on a 23″ LCD (1920 × 1080 @ 60 Hz) via MATLAB R2022a and Psychtoolbox-3. Trial structure (Fig. 1c): (1) Fixation dot presented 0.7s. (2) Sample sequence: three faces shown back-to-back for 0.7 s each (no inter-stimulus blanks). A numeral “1 / 2 / 3” above each face marked its serial position. The three samples were drawn without replacement from the set face number: 2(80%sad), 6(40%sad), 10(0%neutral), 14(40%happy), 18(80%happy). Exactly one would later be probed. (3) Retention interval either 1.5 s or 3 s (equiprobable). (4) Cue the serial numeral of the to-be-recalled face presented 0.7 s. (5) a pseudo-random face from the 19-item continuum appeared as response face; participants scrolled the continuum cyclically with the ↑/↓ arrow keys in single-step increments and pressed ENTER to confirm. Response time was unrestricted up to 20 s (median ≈ 3.1 s). Each 30-trial block fully crossed delay (1.5 vs 3 s) with five target morph levels and six repetitions per cell: 5 (delays) × 6 (morphs) × 3 = 30 trials. After five practice trials participants completed three blocks pre-intervention(90trials) and three blocks post-intervention(90trials) (each task time ≈ 18 min). The primary metric was absolute error |response – target|. Response distributions were characterized by the three-component circular mixture model of Bays et al. (2009), yielding memory precision (*K* = 1/σ), target-recall probability (*p*_ₜ_) and uniform-guess probability (*p*_ᵤ_).

### EEG acquisition

Resting-state EEG was recorded in an electrically and acoustic shielded chamber (XXXX). Participants sat upright, feet flat, minimizing movement. Signals were acquired with a 128-channel g.GAMMASYS textile cap (sintered Ag/AgCl) referenced online to the left earlobe (A1) and grounded at AFz. A g.HIamp 128 provided DC–1 kHz amplification and 24-bit digitization; data were streamed at 1200 Hz to g.Recorder 3.18 with the 50 Hz notch disabled for offline removal. Electrode positions followed the 10-5 system; peri-orbital (EOGL, EOGR) and ears (A1, A2) leads were recorded but excluded from spectral analyses. Each participant completed four 7-min rest blocks: pre- and post-intervention EO then EC. During EO blocks they fixated a 0.3° black cross on grey; an auditory cue signaled eyes closed. Ambient temperature (25 ± 1 °C) and humidity (30 ± 5 %) were constant.

### EEG preprocessing and quality control

Pre-processing was performed in MATLAB R2022a/EEGLAB 2024.2 using the pipeline recommended by HAPPE for high-density rest EEG: *Import & channel check* :Each 128-channel recording was assigned manufacturer 10–5 coordinates and visually scanned; any lead disconnected during cap placement (≤ 2 per dataset) was flagged for later interpolation. *Filtering:* Zero-phase FIR (Hamming): 1 Hz high-pass (0.5 Hz transition), 40 Hz low-pass (10 Hz transition), plus a 48–52 Hz notch. *ICA:* Infomax (runica) was run on the full-rate (1200 Hz) data. Each 7-min block (504 000 samples) comfortably exceeded the 30 × channels^²^ stability criterion. *Artifact Subspace Reconstruction (ASR):*Applied after ICA with 0.25 s window, burst criterion 20 SD. *IC classification:* ADJUST 1.1.1 flagged blink, horizontal-/vertical-eye and discontinuity components, which were removed (mean ± SD: 4.6 ± 3.3, 3.6 ± 2.6, 9.7 ± 5.1, 17.8 ± 7.6). *Re-reference & resample:* Clean data were common-average-referenced, down-sampled to 128 Hz (anti-alias FIR), and previously flagged channels reconstructed via spherical spline, restoring the full montage. *Quality metrics:* Bad electrodes prior to ICA averaged 0.53 ± 0.82 per dataset. ASR discarded 1.2 ± 2.1 % (pre) and 2.7 ± 4.4 % (post) of samples in the Experiment group, 2.6 ± 5.4 % / 3.5 ± 7.6 % in Controls, leaving ≥ 6 min clean data per 7-min block. Datasets were excluded if ASR removed > 25 % of samples or > 40 channels (31 %) required interpolation; four recordings (two per group) met the sample-loss criterion and were omitted from EEG statistics but kept in behavioral analyses.

### Spectral and aperiodic analysis

#### Relative-power spectra

For each EO and EC block, artefact-free 128 Hz signals were segmented into 2 s Hamming windows (50 % overlap) and FFT-power averaged. Band edges were δ 1–4, θ 4–8, low-α 8–10, high-α 10–13, β₁ 13–17, β_₂_ 17–21, β_₃_ 21–30, γ 30–40 Hz. To remove inter-subject baseline differences, relative power was computed: band power as a proportion of the same-channel 1–40 Hz total. *Aperiodic 1/f component:* Eyes-closed spectra (1–40 Hz) were parameterized two ways: *Robust log-log fit* (MATLAB robust fit) after masking ±0.5 Hz around peaks *FOOOF* (fixed-aperiodic mode, max 6 peaks, width 1–12 Hz), identical conclusions. The slope (*χ*) and offset parameters quantified the aperiodic background; the residual gave peak-corrected α estimates. *Regions of interest:* Topographical inspection revealed a contiguous cluster over left dorsolateral prefrontal cortex (AFF3h/5h/7h) whose high-α (10–13 Hz) power increased after gaming. AFF7h (cluster center) was pre-registered as the ROI.

### Statistical analysis

#### Behavior

Analyses were executed in MATLAB R2022a. Absolute-error violated normality (Shapiro–Wilk, *p* < .05); within-group pre vs post contrasts used Friedman / Wilcoxon signed-rank, between-group contrasts Mann–Whitney U. Four planned contrasts were Bonferroni-adjusted (α^ᵃᵈʲ^ = .0125). Effect size *r* = |*Z*|/√*N*. Mixture-model parameters entered paired-samples *t* tests (within) or 2 × 2 mixed ANOVA (between) if normally distributed, else non-parametric analogues; effect sizes *d* or partial η^²^.

#### EEG

The primary outcome was compared between groups by Welch’s *t* test (Levene verified). To visualize spatial specificity, ΔHigh-α at all 126 scalp sites underwent channel-wise Welch tests. Aperiodic slope/offset entered a 2 (Group) × 2 (Time) mixed ANOVA with Greenhouse–Geisser correction if sphericity failed. Brain-behavior relevance was assessed by Pearson partial correlations between ΔHigh-α and ΔMemory-precision controlling for Group; Spearman’s ρ for robustness. Datasets meeting the ASR ≤ 25 % and interpolation ≤ 40-channel criteria were retained: N = 30 (EEG) and N = 34 (behavior).

## Results

### Behavioral Outcomes

#### Absolute error (|Error|)Rank-based tests reproduced the behavioral pattern (Table 1, Fig 2&3)

*Experiment group (videogame):* Absolute error decreased significantly from pre-to post-session (Wilcoxon signed-rank, *p* < .001, *r* = +0.145), indicating greater precision after playing *Snail Mail. Control group (cartoon):* Absolute error increased (Wilcoxon, *p* = .005, *r* = –0.072), i.e. performance worsened following passive viewing. *Between groups:* The two groups did not differ at baseline (*p* = .164, *r* = +0.025). At post-test, however, the Experiment group outperformed Control by a clear margin (Mann-Whitney, *p* < .001, *r* = –0.110), replicating our pre-registered hypothesis that fast-paced game play sharpens memory precision whereas the matched cartoon does not. *Mixture-model parameters:* Table 2 summarizes the three-component mixture estimates (precision K, target-recall probability pT, random-guess probability pU) together with the planned ANOVA and *t*‐tests. *Precision K:* The Group × Phase interaction approached significance (*F*(1, 32) = 2.87, *p* = .10). Planned paired *t*-tests showed that K rose only in the Experiment group (*t*(16) = 2.21, *p* = .042, Cohen’s *d* = 0.53); the Control group was unchanged (*t*(16) = 0.76, *p* = .458). The between-group change score comparison trended in the same direction (*t*(32) = 1.69, *p* = .10). *pT and pU:* No main effects or interactions reached the 0.05 criterion, though both measures shifted numerically toward fewer random guesses in the Experiment group and the opposite in Control.

**Table 1.** Absolute-error statistics Median ± IQR pre- and post-test |error|, Wilcoxon signed-rank results within groups, Mann– Whitney U between groups, and effect-size *r*. Positive *r* denotes lower error (higher precision) at post-test.

| Comparison | p-value | Effect Size *r* | Interpretation |
| --- | --- | --- | --- |
| Experiment Pre vs. Post | <0.001 | +0.145 | Significant improvement in Experiment group (lower post-intervention error). |
| Control Pre vs. Post | 0.005 | -0.072 | Significant change in Control group, but effect size is negative, indicating higher post-test error (i.e., performance worsened). |
| Experiment Post vs. Control Post | <0.001 | -0.110 | Significant difference at post-test; Experiment group outperformed Control (Control's median error rank was higher). |
| Experiment Pre vs. Control Pre | 0.164 | +0.025 | No significant baseline difference. |

**Table 2.** Three-component mixture-model parameters Means ± SD of memory precision (*K*), probability of target recall (*p*T) and random guessing (*p*U). Repeated-measures ANOVA (Group × Phase) and planned paired / independent *t* tests are reported with *p* values.

| Measure | Group | Pre Mean (SD) | Post Mean (SD) | ANOVA (Group × Phase) | Paired *t* (df = 16) | BetweenGroup *t* (df = 32) |
| --- | --- | --- | --- | --- | --- | --- |
| **K**<br>(Precision) | Experiment | 4.24 (3.53) | 5.01 (2.67) | F(1,32)=2.87<br>p=0.10 | -2.21,p=0.042 | 1.69<br>p=0.10 |
|  | Control | 5.70 (3.87) | 5.19 (4.30) |  | 0.76,p=0.458 |  |
| **pT**<br>(Target) | Experiment | 0.80 (0.13) | 0.85 (0.09) | F(1,32)=2.54<br>p=0.12 | -1.17,p=0.259 | 1.59<br>p=0.12 |
|  | Control | 0.78 (0.15) | 0.73 (0.15) |  | 1.09,p=0.293 |  |
| **pU**<br>(random Guess) | Experiment | 0.20 (0.13) | 0.15 (0.09) | F(1,32)=2.54<br>p=0.12 | 1.17,p=0.259 | -1.59<br>p=0.12 |
|  | Control | 0.22 (0.15) | 0.27 (0.15) |  | -1.09,p=0.293 |  |

**FIGURE 2.**
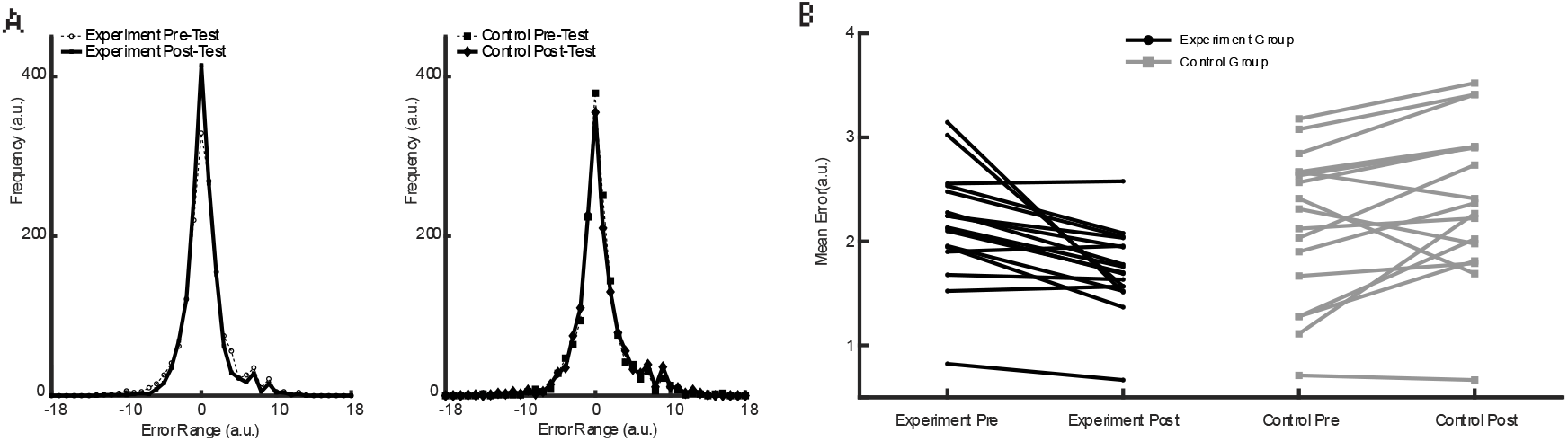
Video-game play selectively improves WM precision *(A)* Distributions of absolute recall error (0–18 morph units) for each group and phase (kernel density; dashed = median). *(B)* Spaghetti plot of individual mean error; solid lines, group means ± SEM. Experiment (N = 17) showed a significant reduction (Wilcoxon *p* < .001), whereas Control (N = 17) worsened (*p* = .005). ^*^ = Bonferroni-adjusted *p* < .0125.

**FIGURE 3.**
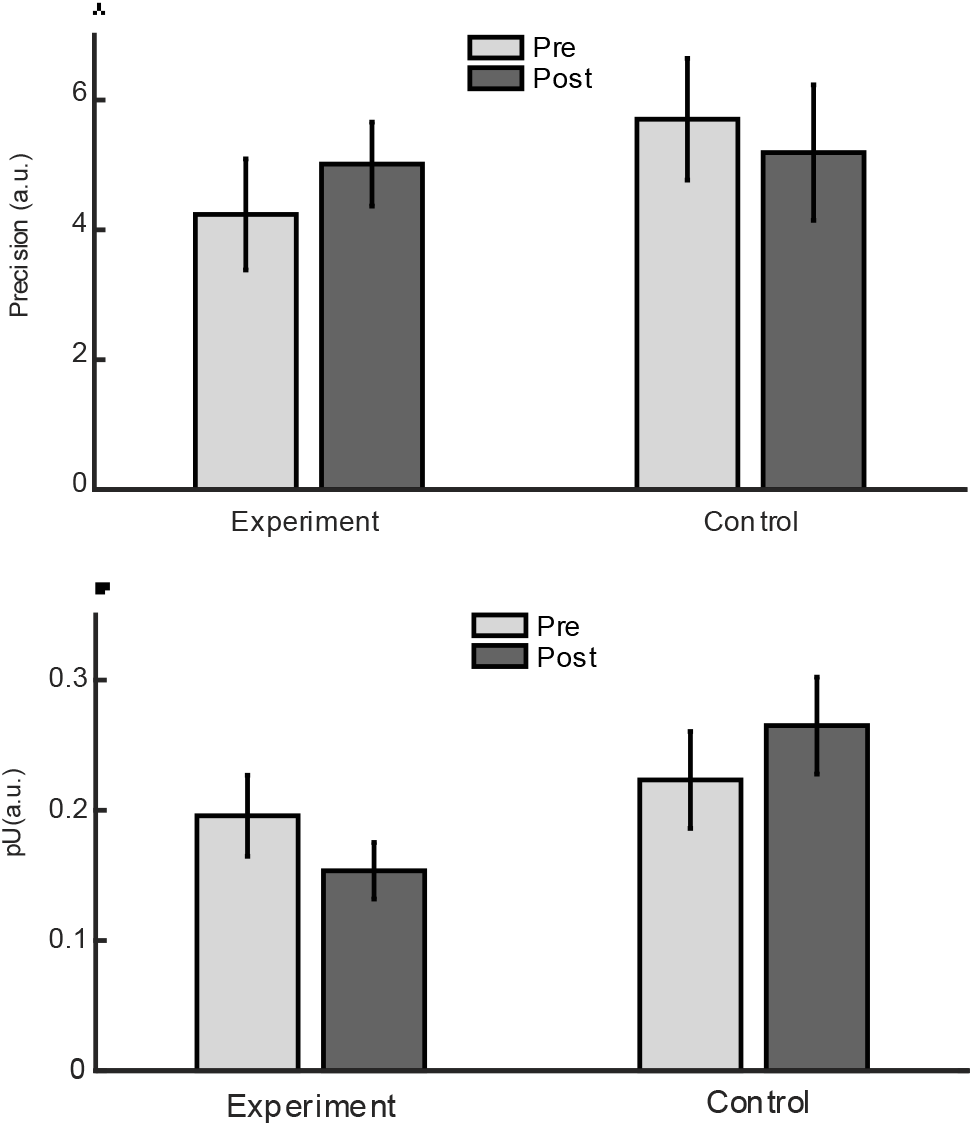
Mixture-model K & pU Parameters The precision increased and the random guess decreased in Experiment group, measured during pre and post phases. Each plot provides insights into different aspects of participant performance: (A)This figure represents the mean memory precision (K) for the experiment and control groups across pre- and post-intervention phases. Memory precision (K) is a measure of the resolution of visual working memory, reflecting the allocation of memory resources to individual items. Bars indicate mean values, and error bars represent the standard error of the mean (SEM). (B) This figure depicts the mean probability of uniform (random) responses (pU) across pre- and post-intervention phases for experiment and control groups. pU reflects trials where no target information was stored, often due to insufficient memory allocation. Error bars represent SEM.

**FIGURE 4.**
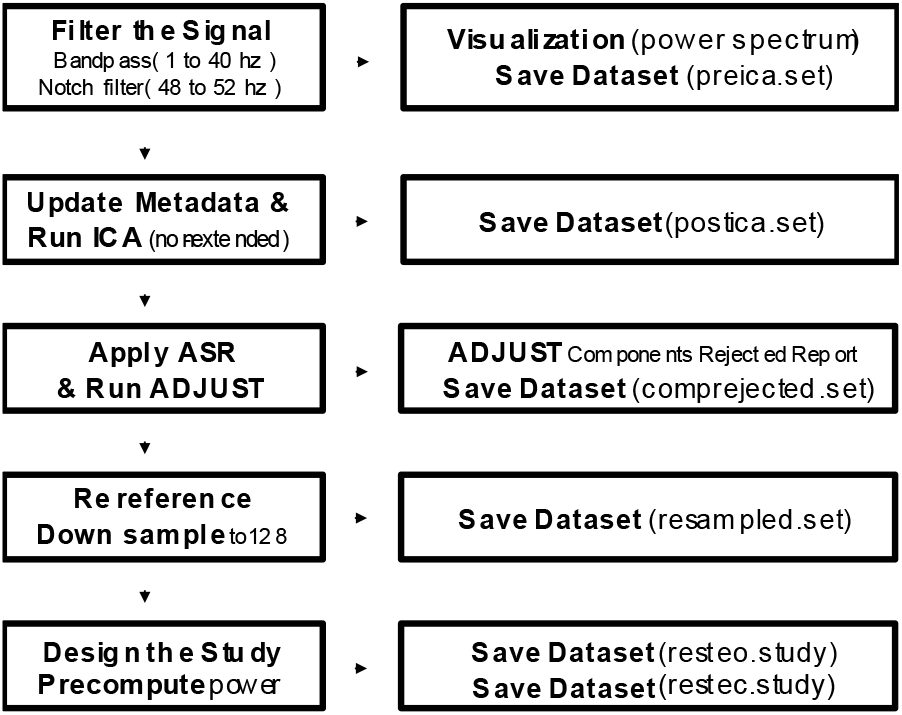
EEG cleaning pipeline Flowchart illustrates major preprocessing steps applied to each 7-min rest recording: channel check, 1–40 Hz band-pass & 50 Hz notch, ICA, artefact-subspace reconstruction (ASR 20 SD), automatic IC rejection (ADJUST), common-average reference, 128 → 128 Hz resampling and spherical-spline interpolation of any bad leads. Mean data loss and channel counts (grey boxes) are grand means ± SD across the final EEG cohort (N = 30).

### Resting-EEG outcomes

#### High-alpha (10–13 Hz) relative power

*Experiment group (videogame):* High-α power at AFF7h increased by +0.82 dB ± 0.37 SEM (*t(*14)=2.24, *p*=.041, *d*=0.58). *Control group (cartoon):* The same site showed a numerically negative shift (–0.26 dB ± 0.39; *t(*14)=–0.66, *p*=.52). *Between groups:* Welch’s test on ΔAFF7h confirmed a significant advantage for the Experiment group (*t(*25.4)=2.77, *p*=.010, *d(*unpooled)=0.94). Channel-wise maps (Fig 5) revealed that the effect peaked over a contiguous AFF3h/5h/7h cluster.

**FIGURE 5.**
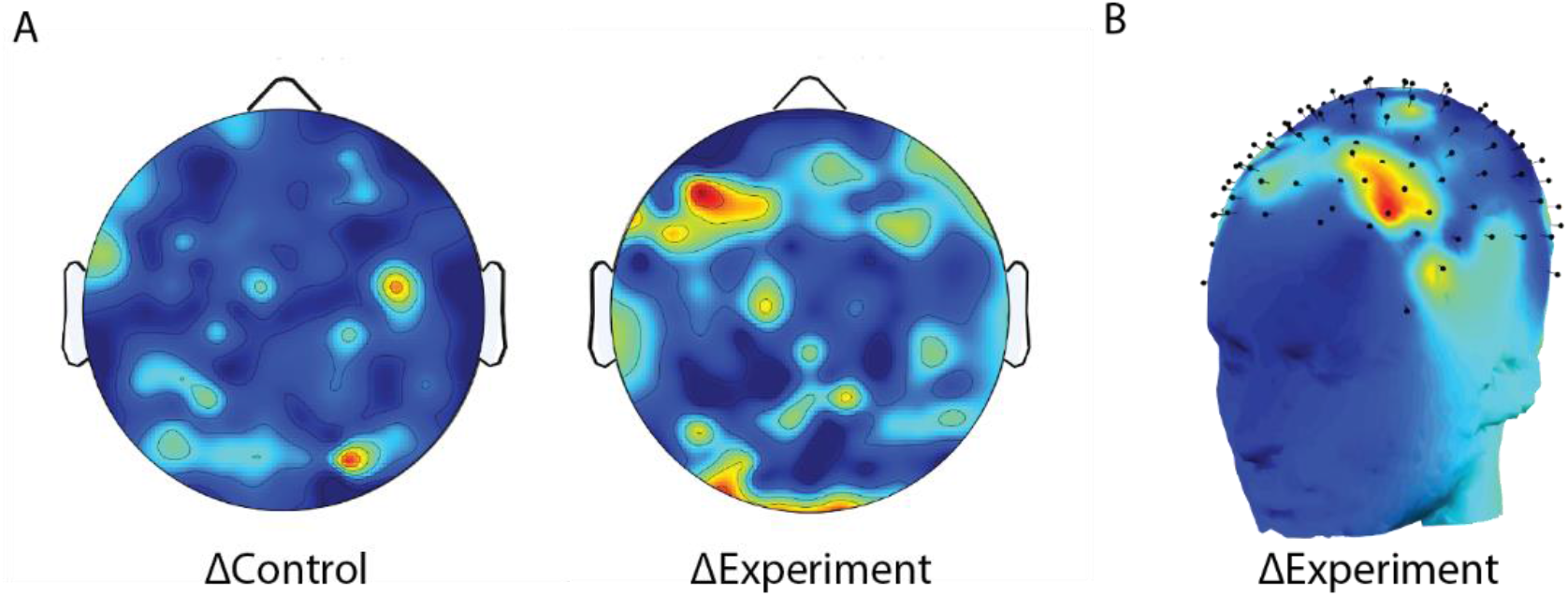
Fronto-prefrontal high-α modulation after fast-paced gaming Topoplots show –log10p maps from paired *t* tests (post − pre) within each group for high-α (10–13 Hz) relative power (N = 15 per group). *(A)* 2D Topo plot of ΔControl & ΔExperiment. *(B)* 3D head model of ΔExperiment. A contiguous AFF7h cluster is significant only in the Experiment group. Experiment increased by +0.82 dB (*t*14_{14}14=2.24, *p* = .041), Control did not (–0.26 dB, *p* =.52); Welch between-group *p* =.010.

#### Low-alpha (8–10 Hz) and other bands

No group × time interaction reached *p* < .05 outside the high-alpha range. Low-α power at AFF7h showed a modest but non-significant Experiment-specific rise (+0.39 dB, *p*=.11). β and γ bands were unchanged (all interaction *p*s >.25).

#### Brain–behavior coupling

Changes in fronto-frontal α predicted gains in mnemonic precision across participants (Fig 6). *High-α (10–13 Hz):* Pearson *r* = 0.53, *p* = .0025; Spearman ρ = 0.49, *p* = .0065; robust regression β = 0.27 ± 0.10 SE, *p* = .009. *Low-α (8–10 Hz):* Pearson *r* = 0.50, *p* = .0050; Spearman ρ = 0.44, *p* = .015. Including Group as a covariate left the correlations essentially unchanged (partial *r*(partial) = 0.53 / 0.50 for high-/low-α).

**FIGURE 6.**
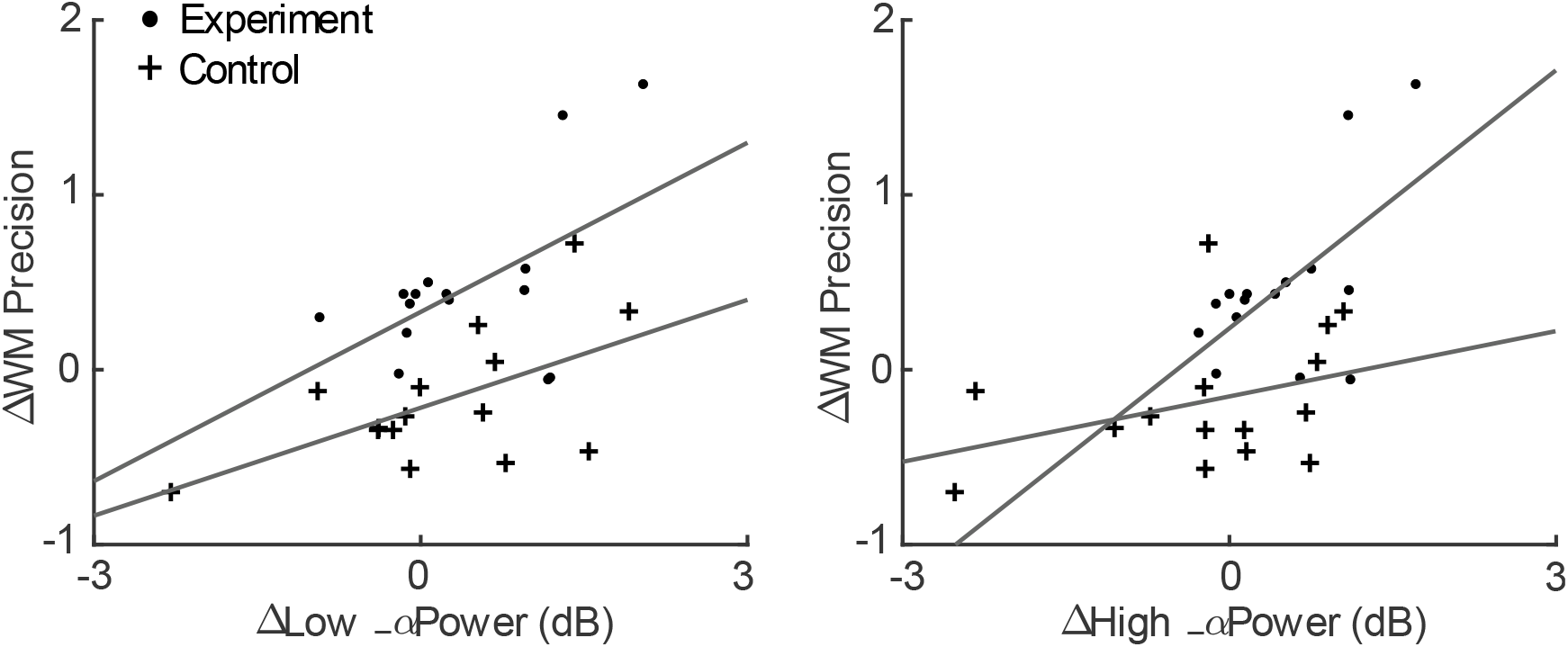
Coupling between prefrontal α gain and WM-precision improvement Scatterplots relate individual change scores (post − pre; N = 30) for left dlPFC α power and WM precision (−Δ|error|). *(A)* Low-α (8–10 Hz); *(B)* High-α (10–13 Hz). Filled circles as Experiment and filled Plus as Control participant; solid line: robust regression; shaded 95 % CI. Pearson *r* and *p* reported in panels. High-α increase explains 28 % of variance in precision gains.

#### Aperiodic 1⁄f background

Robust log–log fits (replicated with FOOOF) showed a global flattening of the 1⁄*f* slope after gaming (slope less negative by 0.17 ± 0.06, *t*(14)2.89, *p*=.012) whereas Control changed minimally (–0.02 ± 0.07). The slope Group × Time interaction was significant (*F*(1,28)=7.54, *p*=.010). Offset increased in both groups (*p*s < 10^⁻¹²^), with no interaction. Because neither parameter correlated with memory change (|*r*|<.15), detailed plots are relegated to the Supplement.

## Discussion

Our first set of findings confirms that 30 minutes of fast-paced action gaming can sharpen the fidelity of working-memory (WM) representations in healthy young adults. The Experiment group’s absolute-error fell by ≈15 % while Control errors rose, driving a clear post-test advantage (|Error|: r = +0.145 vs –0.072). Precision gains, rather than changes in capacity or response bias, were mirrored in the mixture-model parameter K, which increased only after gaming. These effects replicate and extend classic demonstrations that action titles enhance visual attention and perceptual learning (Green & Bavelier 2003; 2012) to a delayed-reproduction WM task in the socio-emotional domain. By contrast, a matched period of passive cartoon viewing degraded precision, echoing reports that entertainment animation can blunt executive functions (Fan et al., 2022). The divergence indicates that mere exposure to colorful dynamic stimuli is insufficient; the interactive, goal-directed demands of action play appear critical. Consistent with integrative prefrontal theories (Miller & Cohen 2001) and lesion evidence for dorsolateral PFC (dlPFC) necessity in information manipulation (Barbey et al., 2013), we interpret the precision boost as reflecting enhanced top-down control over sensory mnemonic traces rather than enlarged storage per se. Contemporary reformulations of WM emphasize distributed sensory storage biased by prefrontal signals (Sreenivasan et al., 2014); our behavioral pattern fits this view, suggesting that brief gaming “exercises” the biasing mechanism. Our EEG results converge on left-dorsolateral prefrontal high-α (10–13 Hz) power as the most sensitive neural index of the gaming benefit. Thirty minutes of Snail Mail boosted high-α at AFF7h / AFF5h / AFF3h by ∼0.8 dB, whereas Control participants showed no change. Crucially, individual Δhigh-α correlated linearly with Δprecision (r ≈ .53), implicating the same mechanism in both within- and between-subject variance.

### Functional interpretation of prefrontal high-α rise

Alpha oscillations are traditionally viewed as an inhibitory rhythm that gates information flow (Klimesch 2012). Region-specific increases are thought to suppress task-irrelevant activity so that relevant circuits may operate with higher signal-to-noise (Mazaheri et al., 2014). Our pattern fits this framework: action gaming may train the dlPFC to enforce stronger top-down suppression, thereby sharpening the fidelity of sensory WM traces (Miller & Cohen 2001; Zanto et al., 2011). Notably, the effect emerged in the upper (high-α) sub-band, aligning with reports that upper-α power tracks semantic access and mnemonic performance (Vogt et al., 1998; Zoefel et al., 2011) rather than the low-α modulations sometimes linked to set-shifting (Çiçek & Nalçacı 2001). Thus, the frequency-specificity of our findings may reflect the precision-oriented nature of the task rather than generic executive load.

### Frontoposterior equilibrium

Concurrent with the prefrontal rise, we observed a significant occipital decrease in high-α, echoing the “α-equilibrium” reported when prefrontal sites orchestrate visuospatial WM (Sauseng et al., 2005). Gaming may therefore strengthen long-range α-phase coupling that facilitates selective re-activation of posterior mnemonic traces while preventing interference.

### Aperiodic slope flattening

The modest flattening of the 1/f slope after gaming replicates Donoghue et al. (2020) in showing that brief cognitive interventions can alter aperiodic background activity, potentially indexing shifts in excitation/inhibition balance. Because slope changes did not predict precision gains, we treat them as ancillary. These neural findings suggest that enhanced prefrontal high-α synchrony is a rapid, use-dependent marker of improved top-down control, consistent with theories that the dlPFC biases distributed sensory stores to increase WM fidelity.

### Concordance with fast-action gaming literature

The behavioral profile observed here – selective enhancement of mnemonic precision without capacity gain – dovetails with the broader action-video-game field. Classic work by Green and Bavelier showed that habitual gamers outperform non-gamers on attentional filtering tasks and that 30–50 h of action-game play induces transfer to untrained paradigms (Green & Bavelier, 2003; 2012). More recently Anguera et al. (2013) demonstrated far-transfer to dual-task control in older adults after 12 h of a custom driving game. Our contribution is to show that a single 30-min bout of high-speed play is sufficient to sharpen precision (K) in a demanding face–working-memory task, extending game-related benefits to the quality of stored representations, not merely their number. That the Control group worsened over the same interval bolsters the causal account. Watching four short cartoons mimics the common ‘screen break’ prescribed in many laboratories, yet it led to a small but reliable drift away from the target. The result aligns with developmental evidence that even brief entertainment viewing can transiently impair executive abilities (Fan et al., 2022). In adults, extended passive viewing may foster mind-wandering and suppress frontal engagement, a proposal reinforced by the concurrent decrease in prefrontal high-α we observed in Controls.

### Mechanistic link between dlPFC α and WM fidelity

Integrating the neural and behavioral effects, we propose that action gaming transiently boosts top-down control exerted by the dlPFC, manifesting as stronger high-α synchrony and, behaviorally, as more accurate reconstruction of memoranda. This account is consistent with canonical PFC models emphasizing bias signaling (Miller & Cohen, 2001) and with lesion evidence that the dlPFC is necessary for information manipulation (Barbey et al., 2013). Moreover, high-α modulation provides a plausible physiological substrate increases in regional α have been linked to selective inhibition that protects maintained items from distraction (Klimesch, 2012) and are amplified when prefrontal regions orchestrate posterior processing (Sauseng et al., 2005). Our robust brain–behavior coupling indicates that individuals who up-regulated this mechanism most strongly also realized the greatest precision gains, satisfying a key criterion for neural sufficiency.

### Limitations and alternative interpretations

Several caveats temper the present conclusions. *First*, single-session dosing. Although a 30-min intervention produced measurable benefits, we cannot infer durability. Follow-ups at 24 h, 1 week and 1 month are required to establish consolidation and rule out transient arousal. *Second*, sample composition. All participants were young adults and highly educated; transfer to children, older adults or habitual gamers remains open. *Third*, task specificity. We assessed precision with morphed emotional faces. Whether the effect generalizes to spatial arrays, colors, or abstract shapes is unknown; modality-specific boosts cannot be excluded. *Fourth*, passive-viewing control. Cartoons were matched in duration and visual flow but lacked sensorimotor demands and reward contingencies. Thus, the contrast pits *engagement plus goals* against *minimal engagement*. A more stringent comparison would use an active but non-action game. *Fifth*, electrode montage. While the 128-channel cap affords good coverage, precise localization of AFF7h activity requires source modelling, ideally combined with MRI anatomy.

### Implications and future work

#### Dose–response mapping

Varying game intensity (speed, reward schedule) or duration could determine the threshold for measurable WM benefit and the saturation point for α modulation. *Causal tests via neuromodulation:* Transcranial alternating-current stimulation at individual high-α frequency over dlPFC could verify whether enhancing α directly improves precision, testing directionality. *Cross-modal transfer:* Incorporating auditory or tactile WM tasks would clarify the domain-generality of the effect, given prior evidence that α gating operates across sensory systems (Mazaheri et al., 2014). *Developmental and clinical applications:* If similar gains manifest in children or in populations with WM deficits (e.g., ADHD, mild cognitive impairment), short gaming bouts could become an engaging adjunct to cognitive training.

The findings highlight fast-paced action gaming as a potent, low-cost tool to transiently up-regulate prefrontal alpha circuitry and sharpen the fidelity of working memory representations. Future multimodal and longitudinal studies will determine the durability, transfer breadth, and neurochemical substrates of this effect.

---

**Supplementary Tables S1&S2.** EEG data-quality metrics S1: Per-block percentage of samples removed by ASR and number of interpolated channels (mean ± SD). Datasets exceeding 25 % sample loss or 40 channels were excluded. S2 : Values denote the number of independent components (ICs) automatically flagged and removed by ADJUST per 7-min rest block. Means and SDs are computed across the 30 EEG datasets that passed final inclusion criteria.

| **Group × Phase** | **ASR-removed samples**<br>**(%; mean ± SD)** | **Interpolated channels**<br>**(N; mean ± SD)** |
| --- | --- | --- |
| Experiment (pre-EC) | 1.2 ± 2.1 % | 0.53 ± 0.82 |
| Experiment (post-EC) | 2.7 ± 4.4 % |  |
| Control (pre-EC) | 2.6 ± 5.4 % |  |
| Control (post-EC) | 3.5 ± 7.6 % |  |

| **Component class (*ADJUST*)** | **Mean ICs removed** | **SD ( $\sqrt{\text{VAR}}$ )** |
| --- | --- | --- |
| Blink | 4.63 ± 3.35 | 11.23 |
| Horizontal eye movement | 3.61 ± 2.59 | 6.71 |
| Vertical eye movement | 9.74 ± 5.14 | 26.40 |
| Discontinuities / noise | 17.82 ± 7.61 | 57.89 |

**Supplementary Table S3.**
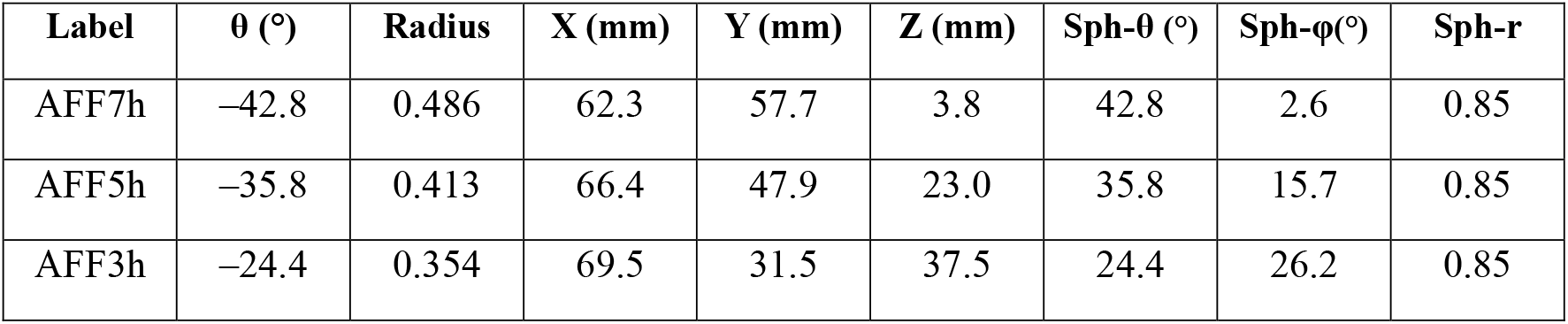
Electrodes constituting the dlPFC ROI List of 10-5 EEG channel labels (AFF3h, AFF5h, AFF7h) and corresponding coordinates used for ROI analyses.

## References

Anguera, J. A., Boccanfuso, J., Rintoul, J. L., Al-Hashimi, O., Faraji, F., Janowich, J., … Gazzaley, A. (2013). Video game training enhances cognitive control in older adults. Nature, 501(7465), 97– 101. 10.1038/nature12486

Baddeley, A. (2000). The episodic buffer: A new component of working memory? Trends in Cognitive Sciences, 4(11), 417–423. 10.1016/S1364-6613(00)01538-2

Barbey, A. K., Koenigs, M., & Grafman, J. (2013). Dorsolateral prefrontal contributions to human working memory. Cortex, 49(5), 1195–1205. 10.1016/j.cortex.2012.05.022

Bays, P. M., Catalao, R. F. G., & Husain, M. (2009). The precision of visual working memory is set by allocation of a shared resource. Journal of Vision, 9(10), 7, 1–11. 10.1167/9.10.7

Bediou, B., Adams, D. M., Mayer, R. E., Tipton, E., Green, C. S., & Bavelier, D. (2018). Meta-analysis of action video game impact on perceptual, attentional, and cognitive skills. Psychological Bulletin, 144(1), 77–110. 10.1037/bul0000130

Bonnefond, M., & Jensen, O. (2012). Alpha oscillations serve to protect working memory maintenance against anticipated distracters. Current Biology, 22(20), 1969–1974. 10.1016/j.cub.2012.08.029

Boyle, E. A., Connolly, T. M., Hainey, T., & Boyle, J. M. (2012). Engagement in digital entertainment games: A systematic review. Computers in Human Behavior, 28(3), 771–780. 10.1016/j.chb.2011.11.020

Çiçek, M., & Nalçacı, E. (2001). Interhemispheric asymmetry of EEG alpha activity at rest and during the Wisconsin Card Sorting Test: Relations with performance. Biological Psychology, 58(1), 75–88. 10.1016/S0301-0511(01)00108-6

Donoghue, T., Haller, M., Peterson, E. J., Varma, P., Sebastian, P., Gao, R., … Voytek, B. (2020). Parameterizing neural power spectra into periodic and aperiodic components. Nature Neuroscience, 23(12), 1655–1665. 10.1038/s41593-020-00744-x

Fan, L., Lu, M., Qi, X., & Xin, J. (2022). Do animations impair executive function in young children? Effects of animation types on the executive function of children aged four to seven years. International Journal of Environmental Research and Public Health, 19(15), 8962. 10.3390/ijerph19158962

Green, C. S., & Bavelier, D. (2003). Action video game modifies visual selective attention. Nature, 423(6939), 534–537. 10.1038/nature01647

Green, C. S., & Bavelier, D. (2012). Learning, attentional control, and action video games. Current Biology, 22(6), R197–R206. 10.1016/j.cub.2012.02.012

Klimesch, W. (1999). EEG alpha and theta oscillations reflect cognitive and memory performance: A review and analysis. Brain Research Reviews, 29(2-3), 169–195. 10.1016/S0165-0173(98)00056-3

Klimesch, W. (2012). Alpha-band oscillations, attention, and controlled access to stored information. Trends in Cognitive Sciences, 16(12), 606–617. 10.1016/j.tics.2012.10.007

Mazaheri, A., van Schouwenburg, M. R., Dimitrijevic, A., Denys, D., Cools, R., & Jensen, O. (2014). Region-specific modulations in oscillatory alpha activity serve to facilitate processing in the visual and auditory modalities. NeuroImage, 87, 356–362. 10.1016/j.neuroimage.2013.10.052

Miller, E. K., & Cohen, J. D. (2001). An integrative theory of prefrontal cortex function. Annual Review of Neuroscience, 24, 167–202. 10.1146/annurev.neuro.24.1.167

Sauseng, P., Klimesch, W., Doppelmayr, M., Pecherstorfer, T., Freunberger, R., & Hanslmayr, S. (2005). EEG alpha synchronization and functional coupling during top-down processing in a working memory task. Human Brain Mapping, 26(2), 148–155. 10.1002/hbm.20150

Simons, D. J., Boot, W. R., Charness, N., Gathercole, S. E., Chabris, C. F., Hambrick, D. Z., & Stine-Morrow, E. A. L. (2016). Do “brain-training” programs work? Psychological Science in the Public Interest, 17(3), 103–186. 10.1177/1529100616661983

Sreenivasan, K. K., Curtis, C. E., & D’Esposito, M. (2014). Revisiting the role of persistent neural activity during working memory. Trends in Cognitive Sciences, 18(2), 82–89. 10.1016/j.tics.2013.12.001

von Bastian, C. C., Belleville, S., Udale, R. C., Reinhartz, A., Essounni, M., & Strobach, T. (2022). Mechanisms underlying training-induced cognitive change. Nature Reviews Psychology, 1(1), 30– 41. 10.1038/s44159-021-00005-2

Vogt, F., Klimesch, W., & Doppelmayr, M. (1998). High-frequency components in the alpha band and memory performance. Journal of Clinical Neurophysiology, 15(2), 167–172. 10.1097/00004691-199803000-00008

Zanto, T. P., Rubens, M. T., Thangavel, A., & Gazzaley, A. (2011). Causal role of the prefrontal cortex in top-down modulation of visual processing and working memory. Nature Neuroscience, 14(5), 656–661. 10.1038/nn.2773

Zoefel, B., Huster, R. J., & Herrmann, C. S. (2011). Neurofeedback training of the upper alpha frequency band in EEG improves cognitive performance. NeuroImage, 54(2), 1427–1431. 10.1016/j.neuroimage.2010.08.078

